# Discovery of a Covalent FEM1B Recruiter for Targeted Protein Degradation Applications

**DOI:** 10.1101/2021.04.15.439993

**Authors:** Nathaniel J. Henning, Andrew G. Manford, Jessica N. Spradlin, Scott M. Brittain, Jeffrey M. McKenna, John A. Tallarico, Markus Schirle, Michael Rape, Daniel K. Nomura

## Abstract

Proteolysis Targeting Chimeras (PROTACs), heterobifunctional compounds that consist of protein-targeting ligands linked to an E3 ligase recruiter, have arisen as a powerful therapeutic modality for targeted protein degradation (TPD). Despite the popularity of TPD approaches in drug discovery, only a small number of E3 ligase recruiters are available for the >600 E3 ligases that exist in human cells. Here, we have discovered a cysteine-reactive covalent ligand, EN106, that targets FEM1B, an E3 ligase recently discovered as the critical component of the cellular response to reductive stress. By targeting Cys186 in FEM1B, EN106 disrupts recognition of the key reductive stress substrate of FEM1B, FNIP1. We further establish that EN106 can be used as a covalent recruiter for FEM1B in TPD applications, in which we demonstrate that a PROTAC linking EN106 to the BET Bromodomain inhibitor JQ1 leads to specific FEM1B- and proteasome-dependent degradation of BRD4 in cells. Our study showcases a covalent ligand that targets a natural E3 ligase-substrate binding site and highlights the utility of covalent ligand screening in expanding the arsenal of E3 ligase recruiters that can be deployed for TPD applications.

## Main Text

Targeted protein degradation (TPD) is based on heterobifunctional compounds, which induce the proximity of E3 ubiquitin ligases to target proteins of interest and thereby elicit their ubiquitination and proteasome-mediated degradation. TPD has become a sought-after therapeutic modality in drug discovery because of its potential ability to specifically eliminate any disease-causing protein in the cell, including classically undruggable targets ^1^. However, a major bottleneck with TPD platforms is the relatively small number of E3 ligase recruiters that are available, despite the >600 E3 ligases that exist in human cells. In fact, most efficient E3 ligase recruiters are based on thalidomide-type immunomodulatory drugs (IMiD) that attract the CUL4 adaptor Cereblon or ligands that recruit the CUL2 adaptor VHL ^2,3^.

Addressing this gap, chemoproteomics-enabled covalent ligand discovery has been deployed to discover E3 ligase recruiters against RNF4, RNF114, DCAF16, and DCAF11 by targeting ligandable cysteines within E3 ligases with covalent ligands and natural products. Ward *et al*. performed a target-based covalent ligand screen using a competitive activity-based protein profiling (ABPP) approach to discover CCW16 that covalently and non-functionally targeted zinc-coordinating cysteines within RNF4. They subsequently demonstrated that linking CCW16 to the BET bromodomain inhibitor JQ1 led to the selective RNF4-dependent degradation of BRD4 ^4^. In a second example, Spradlin *et al*. used ABPP-based chemoproteomic approaches to discover that the natural product nimbolide covalently targeted an N-terminal C8 within a substrate-recognition domain of the E3 ligase RNF114, and that nimbolide could be linked to JQ1 to selectively degrade BRD4 in an RNF114-dependent manner in cancer cells ^5^. Subsequently, Tong, Spradlin *et al*. expanded the utility of nimbolide-based PROTACs demonstrating selective BCR-ABL degradation when linked onto the kinase inhibitor dasatinib ^6^. Luo and Spradlin *et al*. then performed a target-based ABPP screen against RNF114 to discover a fully synthetic covalent RNF114 recruiter EN219 that successfully mimics nimbolide action as an RNF114 recruiter for TPD applications wherein the authors demonstrated RNF114-dependent degradation of BRD4 and BCR-ABL by linking EN219 onto JQ1 or dasatinib, respectively ^7^. In a third successful example, Zhang *et al*. incorporated covalent scout fragments into PROTACs to screen for FKBP12 degradation. Through this screen and using chemoproteomic approaches, they identified a covalent fragment KB02 as a recruiter for the substrate receptor DCAF16 of CUL4-DDB1 E3 ligases for TPD applications ^8^. In a fourth successful example, Zhang *et al*. performed a functional screen with a focused library of electrophilic PROTACs to discover a new recruiter that targets cysteines on DCAF11 ^9^. Thus, these studies have collectively showcased the utility of chemoproteomic approaches and cysteine-targeting ligands to expand the arsenal of E3 ligase recruiters for TPD applications.

The reactivity of Cys residues in cells is maintained by specific signaling pathways that are often centered on E3 ligases and maintain the cellular redox state. Under optimal redox balance, the CUL3 E3 ligase Kelch-like ECH associated protein (KEAP1) sequesters the nuclear transcription factor NRF2 in the cytosol to induce its ubiquitylation and proteasome-mediated degradation. Under conditions of oxidative stress, redox-sensing cysteines on KEAP1 become oxidized to prevent KEAP1-mediated ubiquitination and degradation of NRF2, subsequently allowing NRF2 accumulation and antioxidant gene expression.

Recently, the CUL2 E3 ligase FEM1B was discovered as a critical regulator of the cellular response to persistent depletion of reactive oxygen species (ROS), a condition referred to as reductive stress ^10^. Reductive stress arises from prolonged antioxidant signaling or mitochondrial inactivity and can block stem cell differentiation or lead to diseases, such as cardiomyopathy, diabetes, or cancer. Manford *et al*. discovered that FEM1B, under reductive stress conditions, recognizes reduced cysteines on its substrate FNIP1, leading to FEM1B-dependent FNIP1 ubiquitylation and degradation to restore mitochondrial activity, redox homeostasis and stem cell integrity ^10^. The authors also demonstrated that there was a key cysteine residue C186 that was critical for FEM1B substrate recognition ^10^, suggesting that one could target C186 with a cysteine-reactive covalent ligand to develop a FEM1B recruiter for TPD applications. In this study, we screened a library of cysteine-reactive covalent ligands to discover a covalent FEM1B recruiter and demonstrated that this recruiter can be incorporated into PROTACs to degrade specific protein targets in cells.

To identify a covalent FEM1B recruiter, we screened a library of 566 cysteine-reactive covalent ligands that would inhibit the fluorescence polarization of a TAMRA-conjugated FNIP1^562-591^ degron with recombinant FEM1B **(Fig. 1a, Table S1)**. Through this screen, we identified the chloroacetamide EN106 as the top hit that inhibited FEM1B-FNIP1 degron fluorescence polarization with a 50 % inhibitory concentration (IC50) of 2.2 μM **(Fig. 1b-1c)**. EN106 showed competition against labeling of FEM1B with a cysteine-reactive rhodamine-conjugated iodoacetamide (IA-rhodamine) probe by gel-based ABPP, confirming a direct interaction of EN106 with a cysteine on FEM1B **(Fig. 1d)**. Analysis of EN106 reactivity with recombinant FEM1B by liquid-chromatography-tandem mass spectrometry (LC-MS/MS) analysis of FEM1B tryptic digests revealed an EN106 adduct only on C186—the site that was previously shown to be critical in FEM1B substrate recognition **(Fig. 1e)**^10^.

**Figure 1.**
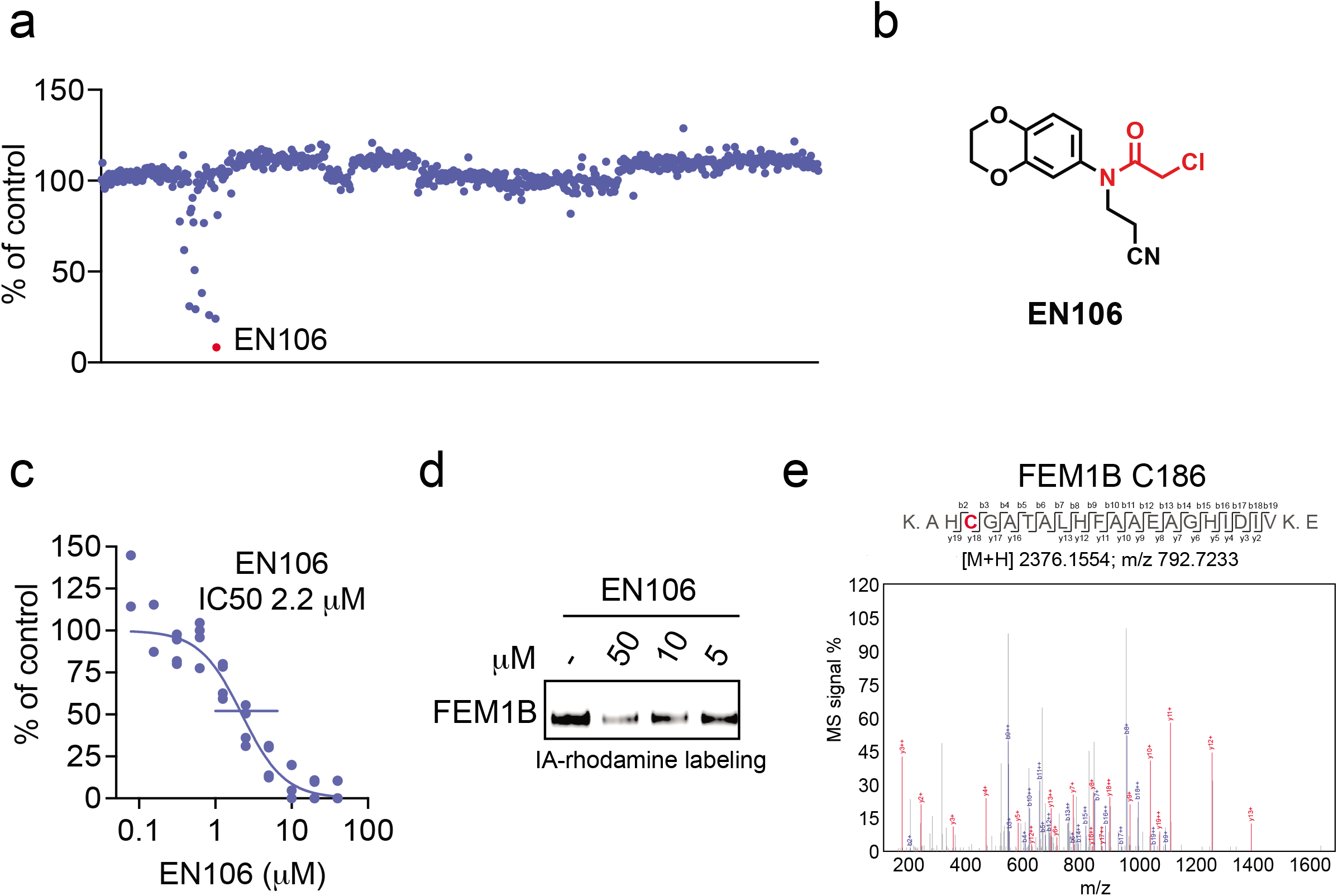
Discovering a FEM1B recruiter. **(a)** Screening a cysteine-reactive covalent ligand library in a fluorescence polarization assay with TAMRA-conjugated FNIP1^562-591^ degron with recombinant MBP-tagged FEM1B. FEM1B was pre-incubated with DMSO vehicle or covalent ligand (50 μM) for 1 hour prior to addition of the TAMRA-conjugated degron. Data for individual compounds screened can be found in **Table S1**. EN106 was the top hit. **(b)** Structure of EN106 with the covalent chloroacetamide handle highlighted in red. **(c)** Dose-response of EN106 inhibition of FEM1B and TAMRA-conjugated FNIP1 interaction assessed by fluorescence polarization expressed as percent fluorescence polarization compared to DMSO vehicle-treated control. **(d)** Gel-based ABPP analysis of EN106. 50 nM pure FEM1B protein was pre-treated with DMSO or EN106 for 30 min at room temperature prior to addition of IA-rhodamine (500 nM, 30 min) at room temperature, after which protein was resolved on SDS/PAGE and visualized by in-gel fluorescence. **(e)** Site of modification of EN106 on FEM1B. FEM1B was labeled with EN106 (50 μM) for 30 min, after which FEM1B tryptic digests were analyzed by LC-MS/MS for the EN106 adduct. Data in **(a)** show average from n=2 biologically independent samples/group. Data in **(c)** shows individual data replicates from n=2-4 biologically independent samples/group. Data in **(d)** shows representative gel from n=3 biologically independent samples/group.

To confirm that EN106 engaged FEM1B in cells, we synthesized NJH-2-030, an alkyne-functionalized derivative of EN106 **(Fig. 2a)**. To maintain engagement of C186, the alkyne was positioned distal to the chloroacetamide by exchanging the benzodioxan for a dihydro[1,4]benzoxazine scaffold. The starting benzoxazine was Boc-protected to give **1** before reduction of the nitro group to provide aniline **2**. Alkylation of **2** with acrylonitrile provided the propionitrile-substituted compound, which was acylated to obtain the chloroacetamide **3**. Boc deprotection and acylation with hex-5-ynoyl chloride provided alkyne probe NJH-2-030 **(Fig. 2a)**.

**Figure 2.**
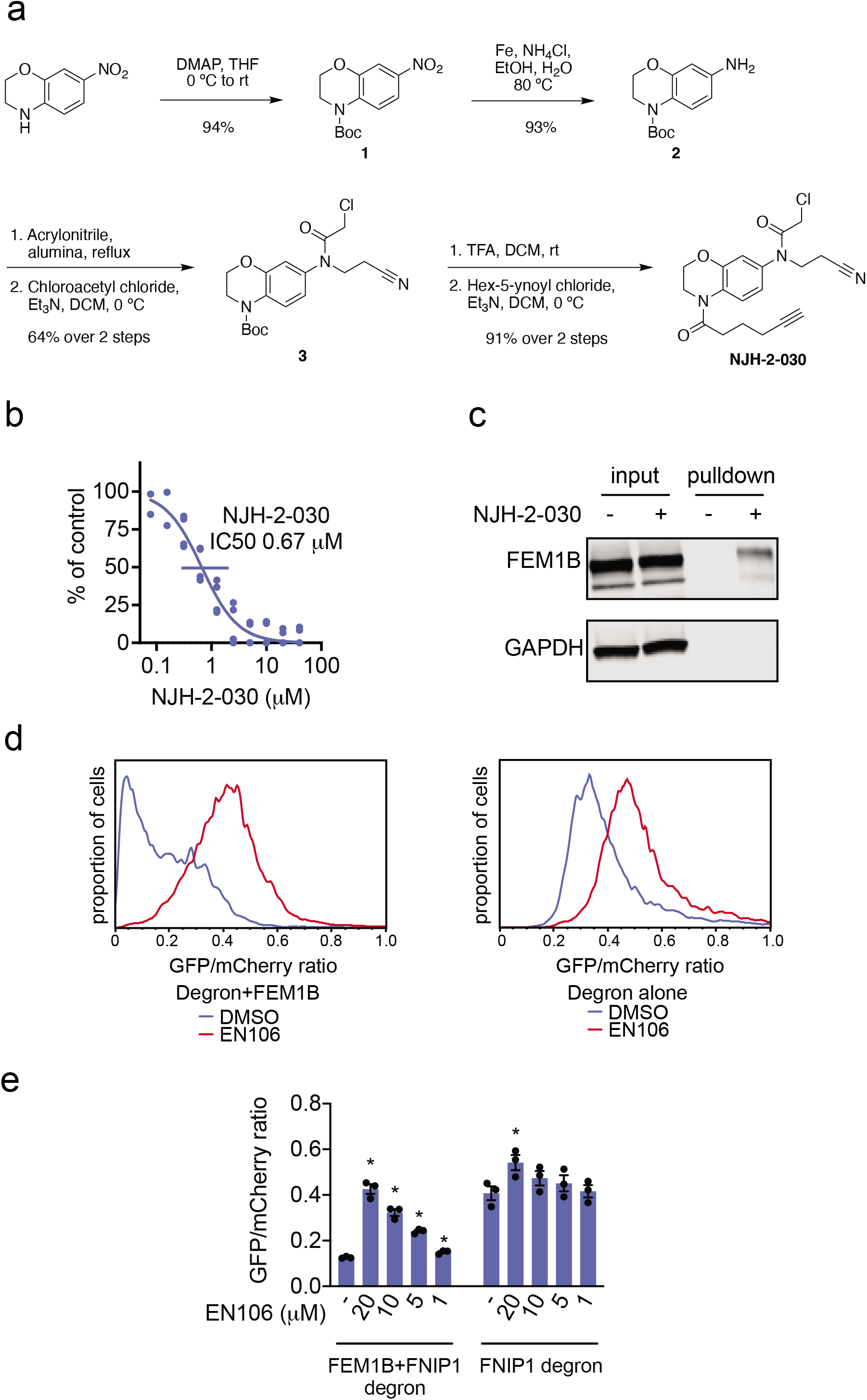
Characterization of EN106 binding with FEM1B. **(a)** Route for synthesis of alkyne-functionalized EN106 probe, NJH-2-030. **(b)** Dose-response of NJH-2-030 inhibition of FEM1B and TAMRA-conjugated FNIP1 interaction assessed by fluorescence polarization expressed as percent fluorescence polarization compared to DMSO vehicle-treated control. **(c)** NJH-2-030 engagement of FEM1B in cells. HEK293T cells were treated with DMSO vehicle or NJH-2-030 (10 μM) for 4 h. Cell lysates were subjected to CuAAC with biotin picolyl azide after which probe-modified proteins were avidin-enriched, eluted, and analyzed by SDS/PAGE and Western blotting for FEM1B or loading control GAPDH. Input levels were also assessed. Shown are representative gels from n=3 biologically independent replicates/group. **(d, e)** Flow cytometry analysis of GFP-FNIP1 degron levels compared to mCherry levels with EN106 treatment in HEK293T cells for 12 h with either basal levels of FEM1B or with transient FEM1B overexpression. Representative flow cytometry traces of DMSO and EN106 (20 μM) treatment groups shown in **(d)** and quantified data of the full dose-response experiment shown in **(e)**. Shown in **(b**,**e)** are individual biological replicate values and average ± sem for n=2-4 biologically independent replicates/group. Significance in **(e)** is expressed as *p<0.05 compared to vehicle-treated controls in each group.

NJH-2-030 maintained inhibitory activity against FEM1B recognition of the FNIP1 degron with an IC50 of 0.67 μM **(Fig. 2b)**. The improved potency with the amide substituent may indicate additional favorable contacts within the FEM1B substrate recognition domain. The NJH-2-030 probe showed FEM1B engagement in HEK293T cells, as demonstrated by FEM1B enrichment from NJH-2-030 treatment in cells, subsequent appendage of biotin-azide by copper-catalyzed azide-alkyne cycloaddition (CuAAC) in cell lysates, avidin-pulldown, and blotting for FEM1B in HEK293T cells compared to vehicle-treated controls **(Fig. 2c)**. An unrelated target GAPDH was not enriched by NJH-2-030 treatment and pulldown **(Fig. 2c)**.

To further demonstrate that EN106 disrupted substrate recognition by FEM1B in cells, we monitored the degradation of GFP linked to a FNIP1 degron compared to IRES driven expression of mCherry from the same plasmid in HEK293T cells by flow cytometry. EN106 treatment significantly stabilized FNIP1 degron-GFP levels, compared to vehicle-treated controls in a dose-responsive manner in FEM1B overexpressing cells **(Fig. 2d)**. EN106 increased FNIP1 reporter levels in cells lacking exogenously expressed FEM1B to a similar extent as previously observed upon deletion of *FEM1B*, indicating that this compound can target the endogenous E3 ligase (**Fig. 2d-2e**). EN106 did not affect the pomalidomide-induced degradation of an unrelated E4F1 degron by the E3 ligase cereblon **(Fig. S1)**. These findings thus indicate that EN106 not only engages, but also inhibits CUL2^FEM1B^ dependent ubiquitylation.

To demonstrate that EN106 could be used as a covalent FEM1B recruiter in TPD applications, we next synthesized NJH-01-106, a compound linking EN106 to the BET bromodomain inhibitor JQ1 that targets BRD4 as well as other BET family proteins **(Fig. 3a)**. Maintaining the core benzoxazine of the alkyne probe NJH-2-030, we first attached an acetate spacer to provide methyl ester **4**. The nitro group was reduced, the resulting aniline **5** mono-alkylated with acrylonitrile, and acylated to provide the chloroacetamide intermediate **6**. The methyl ester was hydrolyzed under mild basic conditions and coupled to amine **7**, a JQ1 derivative, to provide the bifunctional NJH-01-106 **(Fig. 3a)**. NJH-01-106 maintained inhibitory activity against FEM1B recognition of the FNIP1 degron with an IC50 of 1.5 μM **(Fig. 3b)**. NJH-01-106 treatment led to the loss of BRD4 in HEK293T cells with a 50 % degradation concentration (DC50) value of 0.25 μM **(Fig. 3c-3d)**. Loss of BRD4 was attenuated by proteasome and NEDDylation inhibitors, consistent with a proteasome- and Cullin E3 ligase-dependent mechanism of BRD4 degradation **(Fig. 4a-4b)**. BRD4 degradation was also attenuated by pre-treatment of cells with EN106, but not with an unrelated covalent RNF114 E3 ligase recruiter nimbolide **(Fig. 4c)**. In addition, BRD4 degradation was attenuated in FEM1B knockout (KO) cells compared to wild-type (WT) cells, further demonstrating FEM1B-dependent degradation of BRD4 **(Fig. 4d)**. The incomplete rescue we observe in FEM1B KO cells is likely due to the residual FEM1B expression apparent in the FEM1B KO cells or due to other potential off-target E3 ligases. Global proteomic profiling in HEK293T cells treated with NJH-01-106 also showed selective degradation of BRD4 amongst 4446 quantified proteins with only the largely uncharacterized PNMAL1 as an additional off-target **(Fig. 4e; Table S2)**. The observed smaller fold change for BRD4 compared to the Western blot data likely reflect the well-known fold change suppression in TMT-based quantitative proteomics. Together, these results document that covalent modification of a Cys residue in the CUL2 adaptor FEM1B can lead to the development of specific E3 ligase recruiters for targeted protein degradation.

**Figure 3.**
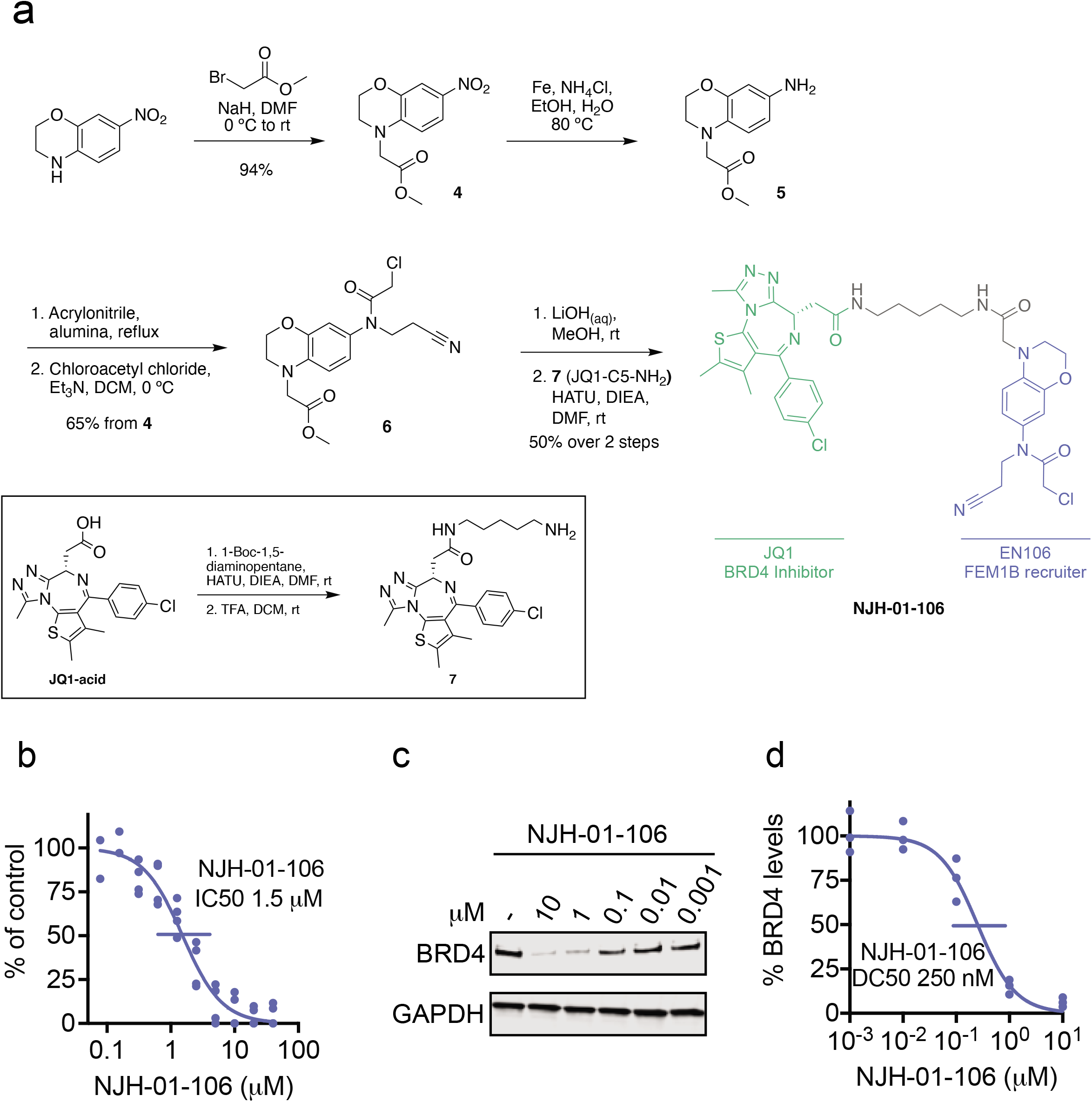
FEM1B-based BRD4 degrader. **(a)** Synthetic route for making FEM1B-based BRD4 degrader NJH-01-106 linking EN106 to JQ1. **(b)** Dose-response of NJH-01-106 inhibition of FEM1B and TAMRA-conjugated FNIP1 interaction assessed by fluorescence polarization expressed as percent fluorescence polarization compared to DMSO vehicle-treated control. **(c)** Degradation of BRD4 by NJH-01-106. NJH-01-106 was treated in HEK293T cells for 8 h and BRD4 and loading control GAPDH levels were detected by Western blotting. **(d)** Quantification of BRD4 degradation from experiment in **(c)** and 50 % degradation concentration value (DC50). Individual biological replicate values shown in **(b)** from n=2-4 biologically independent samples/group. Gel shown in **(c)** is representative of n=3 biologically independent replicates per group which are shown in **(d)**.

**Figure 4.**
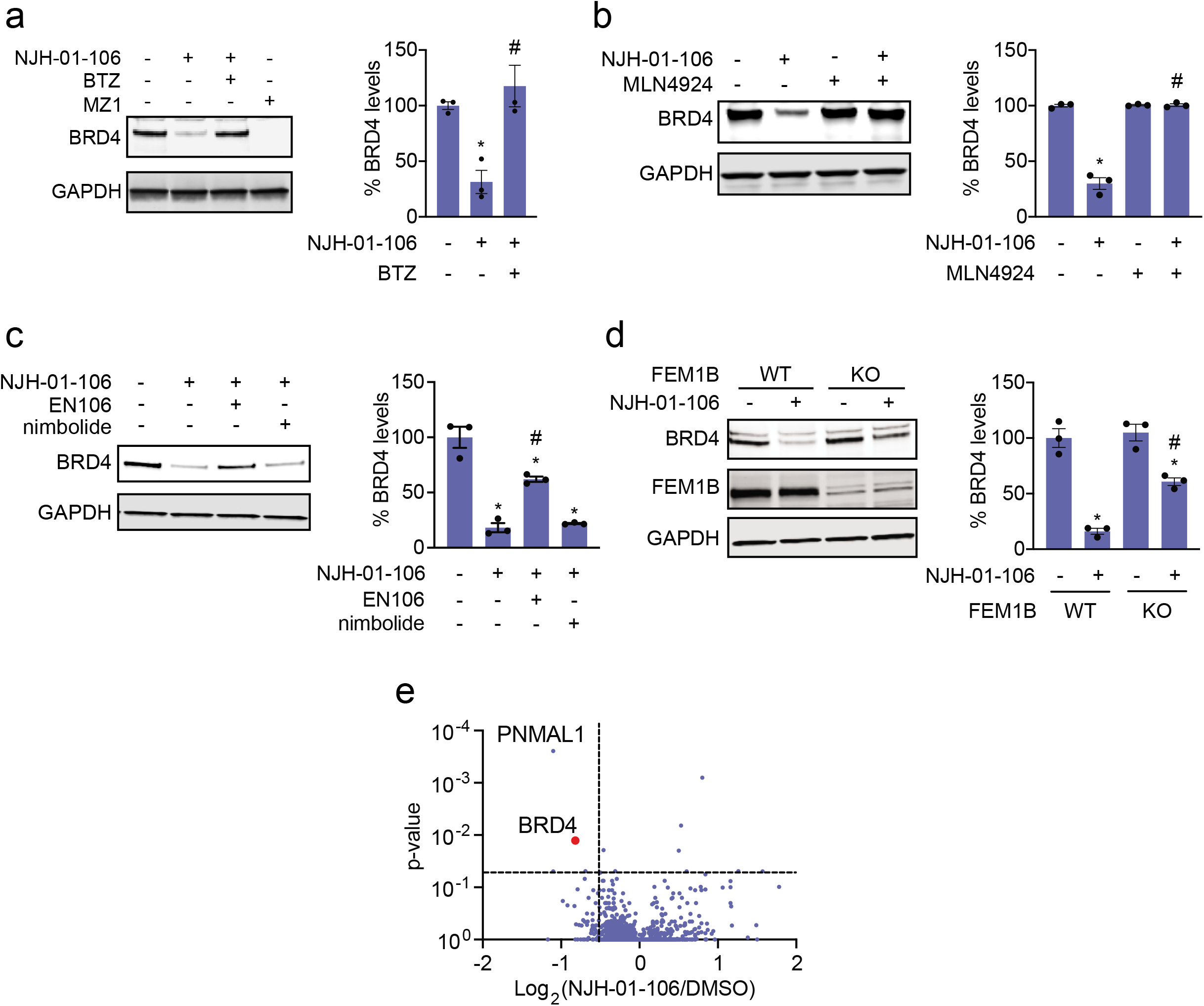
Characterization of FEM1B-based BRD4 degrader. **(a, b)** Proteasome and NEDDylation-dependence of NJH-01-106-mediated degradation of BRD4. HEK293T cells were pre-treated with DMSO vehicle, proteasome inhibitor bortezomib (BTZ) (1 μM), or NEDDylation inhibitor MLN4924 (0.2 μM) for 2 h prior to treatment with DMSO vehicle, NJH-01-106 (10 μM), or MZ1 (1 μM) for 8 h. BRD4 and loading control GAPDH levels were assessed by Western blotting and quantified by Image J. **(c)** Attenuation of BRD4 degradation by EN106, but not nimbolide pre-treatment. HEK293T cells were pre-treated with DMSO vehicle, EN106 (50 μM), or nimbolide (1 μM) for 2 h prior to treatment with DMSO vehicle or NJH-01-106 (10 μM) for 8 h. BRD4 and loading control GAPDH levels were quantified by Image J. **(d)** BRD4 degradation by NJH-01-106 in FEM1B wild-type (WT) and knockout (KO) HEK293T cells. Cells were treated with DMSO vehicle or NJH-01-106 (1 μM) for 8 h. BRD4, FEM1B, and loading control GAPDH levels were assessed by Western blotting and quantified by Image J. **(e)** Proteomic profiling of NJH-01-106 treatment in HEK293T cells. HEK293T cells were treated with DMSO vehicle or NJH-01-106 (1 μM) for 12 h. Protein level changes in cell lysate were quantitatively assessed by TMT-based proteomic profiling. Data shown in **(a-e)** are from n=3 biologically independent samples/group. Gels shown in **(a-d)** are representative gels from n=3 biologically independent samples/group, in which the individual replicate and average ± sem values are shown in bar graphs. Significance is shown as *p<0.05 compared to vehicle-treated groups, and #p<0.05 compared to NJH-01-106-treated groups in **(a-c)** and NJH-01-106-treated WT group in **(d)**.

In this study, we put forth the covalent recruiter EN106 against FEM1B, an E3 ligase involved in reductive stress response, that can be used for TPD applications. While EN106 is an early hit compound with low micromolar potency against FEM1B and requires further medicinal chemistry efforts to improve potency and selectivity, we demonstrate that EN106 targets a Cys residue in FEM1B that is essential for substrate recognition. Moreover, EN106 can be used to develop specific FEM1B recruiters to degrade neo-substrates. Future studies will be needed to determine whether FEM1B-based PROTACs can be used to selectively degrade targets under specific cellular redox conditions. Additionally, determining whether EN106 and more potent derivatives can be used therapeutically to inhibit CUL2^FEM1B^ and disrupt reductive stress signaling through stabilization on FNIP1 in certain cancer settings would be of future interest. Overall, our study underscores the utility of covalent ligand screening in expanding the scope of E3 ligase recruiters for TPD applications.

## Acknowledgement

We thank the members of the Nomura and Rape labs for critical reading of the manuscript. We want to thank Durga Kolla for the E4F1 degron reporter construct. We also want to thank Eddie Wehri and the Henry Wheeler Center for Emerging and Neglected Diseases (CEND) UC Berkeley Drug Discovery Center for providing assistance and equipment for fluorescence polarization assay and screen. We also want to extend our gratitude to the UC Berkeley Cancer Research Laboratory Flow Cytometry Facility. We thank Drs. Hasan Celik, Alicia Lund, and UC Berkeley’s NMR facility in the College of Chemistry (CoC-NMR) for spectroscopic assistance. Instruments in the CoC-NMR are supported in part by NIH S10OD024998. This work was supported by Novartis Institutes for BioMedical Research and the Novartis-Berkeley Center for Proteomics and Chemistry Technologies (NB-CPACT for NJH, SNB, JNS, JMK, JT, MS, DKN), the Mark Foundation for Cancer Research (ASPIRE Award for DKN, NJH, and JNS), the National Institutes of Health (R01CA240981 for DKN), and Howard Hughes Medical Institute (for AGM). MR is an investigator of the Howard Hughes Medical Institute.

## Author Contributions

NJH, AGM, MR, DKN conceived of the project idea, designed experiments, performed experiments, analyzed and interpreted the data, and wrote the paper. SMB, JNS performed experiments, analyzed and interpreted data, and provided intellectual contributions. JMK, MS, JAT provided intellectual contributions to the project.

