## Supporting Information for "Discovery of a Covalent FEM1B Recruiter for Targeted Protein Degradation Applications"

**Supporting Methods**

**Materials**

Cysteine-reactive covalent ligand libraries were purchased from Enamine.

**Fluorescence polarization assay**

Fluorescence polarization assays were performed with purified mouse MBP-FEM1B [1], TAMRA-labeled FNIP1 peptide (5,6-TAMRA- RNKSSLLFKESEETRTPNCNCKYCSHPVLG, Koch Institute/MIT Biopolymers lab). For the screen, 0.5 μl of 2.5 mM compounds were spotted into 384 well non-binding plates (Greiner, 781900). 12.5 μl of 250 nM MBP-FEM1B in binding buffer (40 mM HEPES 7.5, 150 mM NaCl, 0.2 % NP40 substitute, and 100μM TCEP (Tris(2-carboxyethyl)phosphine hydrochloride)) was added to each well and incubated for 1 hour at room temperature. After the incubation, 12.5 μl of 100 nM FNIP1 peptide diluted in binding buffer was added bringing the final concentration to 50 nM for the peptide and 125 nM for MBP-FEM1B. After 1 hour of incubation plates were measured on a Perkin Elmer 2104 Envision plate reader. Data was calculated from mP values (1000*(S-G*P)/(S+G*P), S = 595s channel 2 and P = 595p channel 1, G=1.1) subtracted from peptide only plate. Dose response assays were performed as above, but with 250 nM MBP-FEM1B treated with indicated concentrations of compound (relative to the final reaction volume) or DMSO in separate tubes for 1 hour at room temperature. 12.5 μl of treated MBP-FEM1B (125 nM final) was then added to 12.5 μl of peptide (10 nM final) and incubated with gentle rocking for 30 minutes before measuring fluorescence polarization.

**Gel-Based ABPP**

Recombinant MBP-FEM1B$^{1-377}$ (0.1μg/sample) was pre-treated with either DMSO vehicle or EN106 or at 37℃ for 30 min in 25 μL of PBS, and subsequently treated with of IA-Rhodamine (concentrations designated in figure legends) (Setareh Biotech) at room temperature for 1 h. The reaction was stopped by addition of 4×reducing Laemmli SDS sample loading buffer (Alfa Aesar). After boiling at 95℃ for 5 min, the samples were separated on precast 4−20% Criterion TGX gels (Bio-Rad). Probe-labeled proteins were analyzed by in-gel fluorescence using a ChemiDoc MP (Bio-Rad).

**Cell Culture**

HEK293T cells were obtained from the UC Berkeley Cell Culture Facility and cultured in DMEM (Gibco) containing 10% (v/v) fetal bovine serum (FBS) and maintained at 37 °C with 5% $CO_2$. The FEM1B knockout HEK293T cell line was generated as described by Manford et al [1].

**NJH-2-030 Pulldown and Blotting for FEM1B**

HEK293T cells were treated with DMSO or 10 μM NJH-2-030 *in situ* for 8h. Cells were harvested, lysed via sonication and normalized to 2.0 mg/mL. Following normalization, 100 μL of each lysate sample was removed for Western blot analysis of input, and 500 μL of each lysate sample was incubated for 1 h at room temperature with 10 μL of 5 mM biotin picolyl azide (in water) (Sigma Aldrich 900912), 10 μL of 50 mM TCEP (in water), 30 μL of TBTA ligand (0.9 mg/mL in DMSO:t-butanol = 1:4), and 10 μL of 50 mM Copper (II) Sulfate (12.5 mg/mL in water). Proteins were precipitated, washed 3 x 1 mL with cold MeOH, resolubilized in 1 mL of 1.2% SDS/PBS (w/v), heated for 5 min at 90 °C, and centrifuged to remove any insoluble components. 1 mL of each resolubilized sample was then transferred to 15 mL conical tubes containing 5 mL PBS with 85 μL streptavidin resin (Thermo Scientific 53114) to give a final SDS concentration of 0.2%. Samples were incubated with the streptavidin beads at 4 °C overnight on a rotator. The following day the samples were warmed to room temperature and washed with 0.2% SDS, then transferred to spin columns and further washed 3 x with 500 μL PBS and 3 x with 500 μL water to remove non-probe-labeled proteins. The washed beads were resuspended in 100 μL PBS, transferred to 1.5 mL eppendorf low-adhesion tubes, combined with 30 μL Laemmli Sample Buffer (4 x) and heated to 95 °C. Proteins in each sample were then analyzed by Western blotting to look for enriched FEM1B versus non-enriched GAPDH.

**Flow Cytometry Analysis of GFP-FNIP1 degron/mCherry**

HEK293T cells/well were seeded into 6 well plates. The next day the cells were transfected with 0.1 μg of pCS2-GFP-FNIP1$^{562\text{-}591}$-IRES-mCherry or 0.1 μg of pCS2-E4F1$^{23432}$-GFP-IRES-mCherry, with 0.075 μg pCS2-3xFLAG-FEM1B as indicated. Empty pCS2 was added to 2 μg total DNA for each transfection in 300 μl Opti-MEM (Thermo Fisher, 31985-070) with 12 μg polyethyleneimine (PEI, Polysciences 23966-1). Each well was transfected with 65 μl of the transfection mix. 12 hours post-transfection, indicated concentrations of EN106 or

DMSO was added. After 12 hours of EN106 treatment, cells were trypsinized, spun down, resuspended in DMEM + 10% FBS and analyzed on Fortessa X20. Data was processed using FlowJo and all quantifications are the median GFP/mCherry ratios. For the pomalidomide treated cells, 10 μM pomalidomide (MedChemExpress, HY-10984) was added for 4 hours before analyzing.

**Cell Lysis Protocol**

Pelleted cells were lysed with RIPA lysis buffer (50mM Tris-HCl, 165mM NaCl, 12mM sodium deoxycholate, 1% Triton X-100, 0.01% SDS), protein concentration normalized using a BCA assay (Thermo).

**Western Blot Protocol**

Proteins were resolved by SDS–PAGE (4–20% TGX gels, Bio-Rad Laboratories, Inc.) and transferred to nitrocellulose membranes using the Trans-Blot Turbo transfer system (Bio-Rad). Membranes were blocked with 5% BSA in Tris-buffered saline containing Tween 20 (TBST) solution for 1 h at room temperature, washed in TBST and probed with primary antibody diluted in diluent, as recommended by the various manufacturers, overnight at 4 °C. Primary antibodies used were: BRD4 (Cell Signaling Technologies #13440), GAPDH (ProteinTech 60004-1-Ig), FEM1B (ProteinTech 19544-1-AP). Following washes with TBST, the blots were incubated in the dark with secondary antibodies purchased from Li-Cor Biosciences and used at 1:10,000 dilution in 5% BSA in TBST at room temperature for 1 h. Blots were visualized using an Odyssey Li-Cor scanner after additional washes. Protein intensity was quantified using ImageJ software.

**Quantitative TMT Proteomics Analysis**

Quantitative TMT-based proteomic analysis was performed as previously described [2]. Acquired MS data was processed using Proteome Discoverer v. 2.2.0.388 software (Thermo) utilizing Mascot v 2.5.1 search engine (Matrix Science, London, UK) together with Percolator validation node for peptide-spectral match filtering [3]. Data was searched against Uniprot protein database (canonical human and mouse sequences, EBI, Cambridge, UK) supplemented with sequences of common contaminants. Peptide search tolerances were set to 10 ppm for precursors, and 0.8 Da for fragments. Trypsin cleavage specificity (cleavage at K, R except if followed by P) allowed for up to 2 missed cleavages. Carbamidomethylation of cysteine was set as a fixed modification, methionine oxidation, and TMT-modification of N-termini and lysine residues were set as variable modifications. Data validation of peptide and protein identifications was done at the level of the complete dataset consisting of combined Mascot search results for all individual samples per experiment via the Percolator validation node in Proteome Discoverer. Reporter ion ratio calculations were performed using summed abundances with most confident centroid selected from 20 ppm window. Only peptide-to-spectrum matches that are unique assignments to a given identified protein within the total dataset are considered for protein quantitation. High confidence protein identifications were reported using a Percolator estimated <1% false discovery rate (FDR) cut-off. Differential abundance significance was estimated using a background-based ANOVA with Benjamini-Hochberg correction to determine adjusted p-values.

**Data Availability Statement**

The datasets generated during and/or analyzed during the current study are available from the corresponding author on reasonable request.

**Code Availability Statement**

Data processing and statistical analysis algorithms from our lab can be found on our lab's Github site: https://github.com/NomuraRG, and we can make any further code from this study available at reasonable request.

**Synthetic Characterization and Methods**

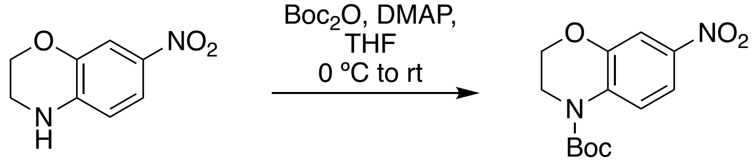


1

tert-butyl 7-nitro-2,3-dihydro-4H-benzo[b][1,4]oxazine-4-carboxylate (**1**): 7-nitro-3,4-dihydro-2H-benzo[b][1,4]oxazine (1.0 g, 5.55 mmol) was dissolved in THF (20 mL) and DMAP (67 mg, 0.55 mmol) was added at 0 ºC followed by $Boc_2O$ (1.45 g, 6.66 mmol). The ice bath was removed after 5 minutes and the mixture stirred overnight at room temperature. The reaction was basified with 15 mL 1M NaOH, stirred for 2 hours, concentrated, and the aqueous mixture extracted with EtOAc. Organic extracts were washed with 1M HCl, brine, dried over $Na_2SO_4$, and concentrated to provide **1** as an orange solid (1.46 g, 5.21 mmol, 94%). LCMS $[M+H]^+$ calc 281.11, found 281.1. 1H NMR (300 MHz, CDCl3) δ 8.11 (d, J = 9.1 Hz, 1H), 7.81 (s, 2H), 4.35 (s, 2H), 3.96 (s, 2H), 1.61 (s, 9H).

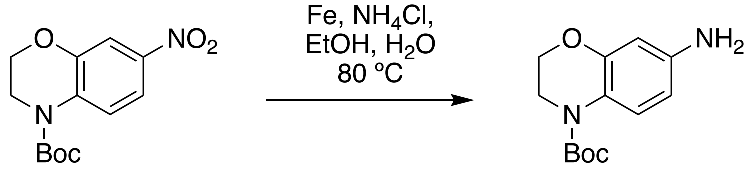


1 2

tert-butyl 7-amino-2,3-dihydro-4H-benzo[b][1,4]oxazine-4-carboxylate (**2**): Nitrobenzoxazine 1 (1.46g, 5.21mmol) was dissolved in EtOH (20mL) and water (5mL) before NH4CL (1.78g, 33.3 mmol) was added. The mixture was heated to 60 ºC, Fe powder (932mg, 16.7 mmol) was added, and the mixture heated at 80 ºC for 13 hours. The reaction was filtered through Celite, and celite washed with EtOAc. Water was added to the filtrate and the mixture extracted with EtOAc. Combined organic extracts were washed with brine, dried over $Na_2SO_4$, and concentrated to provide the aniline **2** as an orange oil (1.21g, 4.82 mmol, 93%). LCMS $[M+H]^+$ calc 251.1, found 251.1. 1H NMR (300 MHz, CDCl3) δ 7.54 (s, 1H), 6.33 – 6.21 (m, 2H), 4.24 (s, 2H), 3.85 (s, 2H), 3.58 (s, 2H), 1.56 (s, 9H).

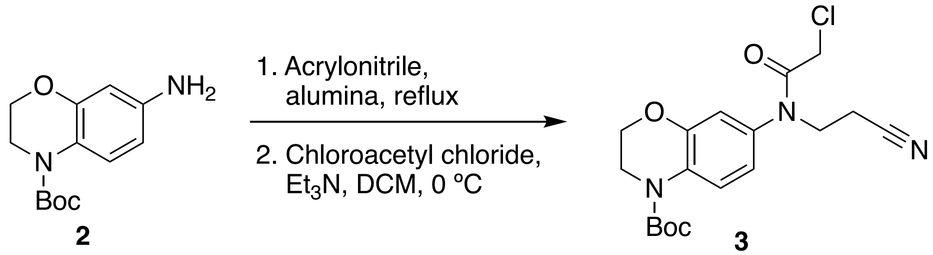


2 3

tert-butyl 7-(2-chloro-N-(2-cyanoethyl)acetamido)-2,3-dihydro-4H-benzo[b][1,4]oxazine-4-carboxylate (**3**): Aniline 2 (1.41 g, 5.66 mmol) was dissolved in acrylonitrile (20 mL). Alumina (1.13 g, 11.1 mmol) was added and the mixture refluxed for 48 hours, before being diluted with EtOAc and filtered through Celite to remove alumina. The filtrate was concentrated to provide the alkylated aniline intermediate (1.82 g) as an orange oil, which was as a mixture of double and single alkylation. LCMS $[M+H]^+$ calc 304.2, found 304.2. One sixth of this crude intermediate, (304 mg, ~1.0 mmol) was dissolved in DCM (6 mL), the solution cooled to 0 ºC, and TEA (556 mL, 4 mmol) was added followed by chloroacetyl chloride (202 mL, 2.5 mmol). The solution was stirred at 0 ºC for 5 minutes and allowed to warm to rt over 1 hour. Aqueous $NaHCO_3$ was added, the mixture partitioned, the aqueous layer extracted with DCM. Combined organic extracts were dried over $Na_2SO_4$, concentrated and purified by silica gel chromatography to obtain chloroacetamide **3** (244 mg, 0.64 mmol, 68% over two steps) as an orange solid. LCMS $[M+H]^+$ calc 380.1, found 308.1. 1H NMR (400 MHz, CDCl3) δ 8.01 (s, 1H), 6.87 – 6.79 (m, 2H), 4.35 – 4.28 (m, 2H), 4.06 – 3.89 (m, 6H), 2.75 (t, J = 6.8 Hz, 2H), 1.60 (s, 9H).

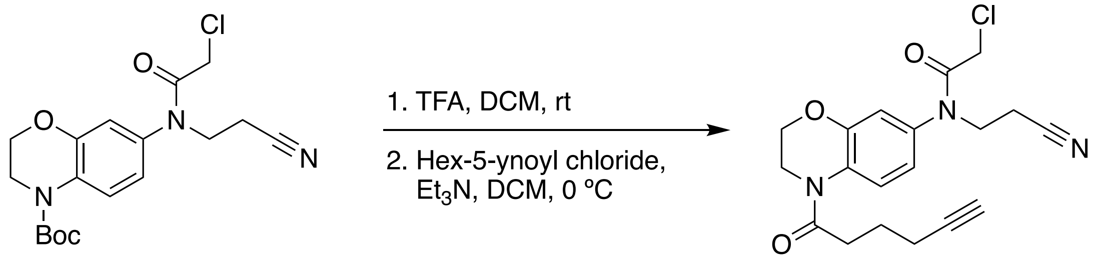

**3** **NJH-2-030**

2-chloro-N-(2-cyanoethyl)-N-(4-(hex-5-ynoyl)-3,4-dihydro-2H-benzo[b][1,4]oxazin-7-yl)acetamide (**NJH-2-030**): Oxalyl chloride (150 mL of 2.0 M solution in DCM, 0.3 mmol) was added to solution of 5-hexynoic acid (17 mL, 0.15 mmol) and 1 drop of DMF in 1 mL in DCM at room temperature. The solution was stirred for 30 minutes before being concentrated to give crude hex-5-ynoyl chloride as a pink foam. Meanwhile, benzoxazine **3** (20 mg, 0.052 mmol) was dissolved in DCM (400 mL) and TFA (400 mL) was added. The solution turned light purple and after stirring 5 min at rt the mixture was concentrated under vacuum, and the resulting aniline was redissolved in DCM (0.5 mL). At 0C, TEA (109 mL, 0.78 mmol) was added to the solution followed by the crude hex-5-ynoyl chloride dissolved in DCM (1 mL). The reaction was stirred for 5 minutes at 0 ºC, water was added, the mixture partitioned and the aqueous layer extracted with DCM. Combined organic extracts were dried over $Na_2SO_4$, concentrated, and purified by silica gel chromatography to obtain **NJH-2-030** (17 mg, 0.047 mmol, 91%). HRMS $[M+H]^+$ calc 374.1193, found 374.1247. 1H NMR (400 MHz, DMSO) δ 7.94 (s, 1H), 7.03 (d, J = 2.5 Hz, 1H), 6.93 (d, J = 8.7 Hz, 1H), 4.30 (t, J = 4.5 Hz, 2H), 4.07 (s, 2H), 3.94 – 3.82 (m, 4H), 2.80 (s, 1H), 2.75 – 2.64 (m, 4H), 2.27 – 2.19 (m, 2H), 1.76 (p, J = 7.2 Hz, 2H). 13C NMR (151 MHz, CDCl3) δ 170.93, 166.76, 147.68, 127.10, 125.66, 119.46, 117.48, 116.71, 83.31, 69.48, 60.39, 46.18, 41.62, 32.78, 23.77, 21.05, 17.76, 16.38, 14.20.

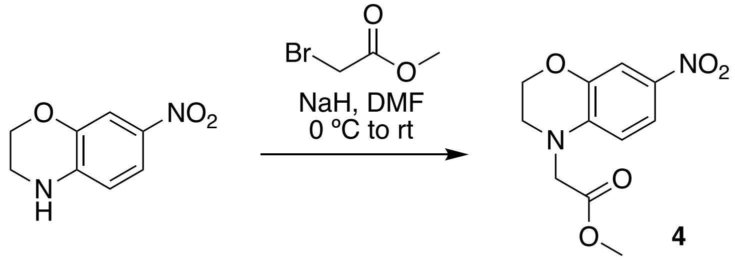

**4**

methyl 2-(7-nitro-2,3-dihydro-4H-benzo[b][1,4]oxazin-4-yl)acetate (**4**): 7-nitro-3,4-dihydro-2H-benzo[b][1,4]oxazine (1.0 g, 5.55 mmol) was dissolved in DMF (20 mL) and cooled to 0 ºC. NaH (233 mg, 5.83 mmol, 60% in mineral oil) was then added to the solution portionwise, and allowed to stir at 0 ºC for 30 minutes before methyl bromoacetate (650 mL, 4.43 mmol) was added dropwise. The solution was then allowed to warm to room temperature and stirred for 2 hours, before it was again cooled to 0 ºC and diluted with water (80 mL). The resulting suspension was filtered to obtain **4** (1.32 g, 5.23 mmol, 94%) as a bright yellow powder. LCMS $[M+H]^+$ calc 253.07, found 253.2. 1H NMR (300 MHz, CDCl3) δ 7.83 (d, J = 9.2 Hz, 1H), 7.75 (s, 1H), 6.50 (d, J = 8.8 Hz, 1H), 4.34 (s, 2H), 4.17 (d, J = 1.9 Hz, 2H), 3.81 (s, 3H), 3.62 (s, 2H).

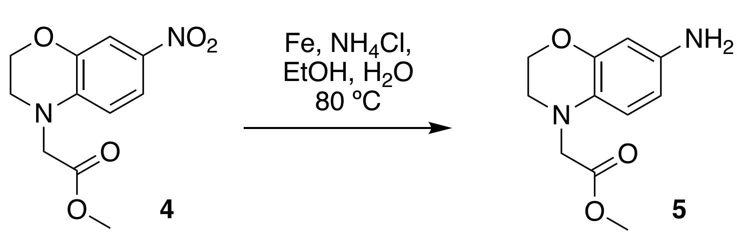

**4** **5**

methyl 2-(7-amino-2,3-dihydro-4H-benzo[b][1,4]oxazin-4-yl)acetate (**5**): Nitrobenzoxazine **4** (1.32 g, 5.23 mmol) was dissolved in EtOH (30 mL) and water (8 mL) before $NH_4CL$ (1.68 g, 31.4 mmol) was added. The mixture was heated to 60 ºC, Fe powder (876 mg, 15.7 mmol) was added, and the mixture heated at 80 ºC for 17 hours. The reaction mixture was filtered through Celite, and the celite pad washed with EtOAc. Water was added to the filtrate and the mixture extracted with EtOAc. Combined organic extracts were washed with brine, dried over $Na_2SO_4$, and concentrated to provide **5** (270mg, 1.0 mmol, 102%) as a brown oil without further purification. LCMS $[M+H]^+$ calc 223.1, found 223.1. 1H NMR (300 MHz, DMSO) δ 6.30 (d, J = 8.3 Hz, 1H), 6.09

– 6.03 (m, 2H), 4.49 (d, J = 15.6 Hz, 2H), 4.15 (s, 2H), 4.05 (d, J = 10.8 Hz, 2H), 3.63 (d, J = 2.0 Hz, 3H), 3.37 (s, 2H).

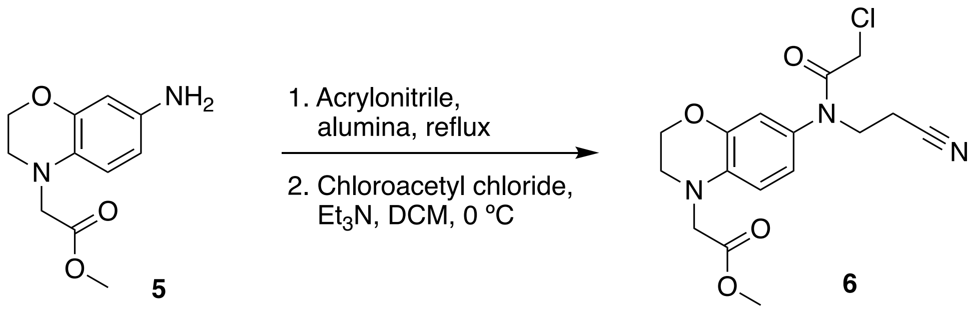


methyl 2-(7-(2-chloro-N-(2-cyanoethyl)acetamido)-2,3-dihydro-4H-benzo[b][1,4]oxazin-4-yl)acetate (**6**): Aniline **5** (974 mg, 4.20 mmol) was dissolved in acrylonitrile (15 mL) and basic alumina (857 mg, 8.40 mmol) was added and the mixture stirred at 80 ºC for 36 hours. The reaction mixture was then diluted with EtOAc and filtered through Celite. Water was added to the filtrate, the mixture partitioned, and the aqueous layer extracted with EtOAc. Combined organic extracts were washed with brine, dried over $Na_2SO_4$, concentrated, and the crude residue purified by silica gel chromatography to obtain a mixture of double and single alkylated aniline (886 mg) as an amber oil. LCMS [M+H]+ calc 276.13, found 276.1. One fourth of this intermediate (215 mg, ~0.78 mmol) was dissolved in DCM (4 mL). The solution was cooled to 0C and TEA (326 mL, 2.34 mmol) was added, followed by chloroacetyl chloride (92 mL, 1.17 mmol). After 15 minutes at 0 ºC, the reaction mixture was concentrated and the crude residue purified by silica gel chromatography to obtain chloroacetamide **6** (229 mg, 0.65 mmol, 62% over two steps) as amber oil. LCMS $[M+H]^+$ calc 352.1, found 352.1. 1H NMR (400 MHz, DMSO) δ 6.82 – 6.72 (m, 2H), 6.62 (dd, J = 8.5, 2.1 Hz, 1H), 4.29 – 4.18 (m, 4H), 4.08 – 3.98 (m, 3H), 3.80 (t, J = 6.7 Hz, 2H), 3.66 (d, J = 2.2 Hz, 3H), 3.44 (s, 2H), 2.68 (t, J = 6.7 Hz, 2H).

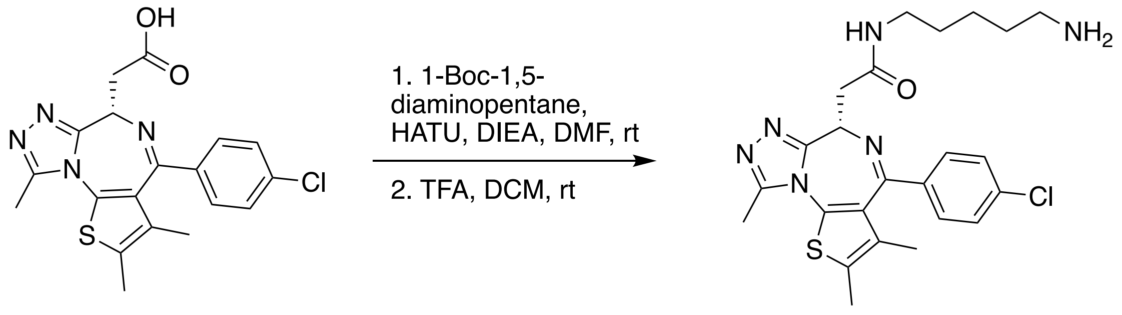


**JQ1-acid**

**7**

(S)-N-(5-aminopentyl)-2-(4-(4-chlorophenyl)-2,3,9-trimethyl-6H-thieno[3,2-f][1,2,4]triazolo[4,3-a][1,4]diazepin-6-yl)acetamide (**7**) was prepared as described by Koblan et al. [4].

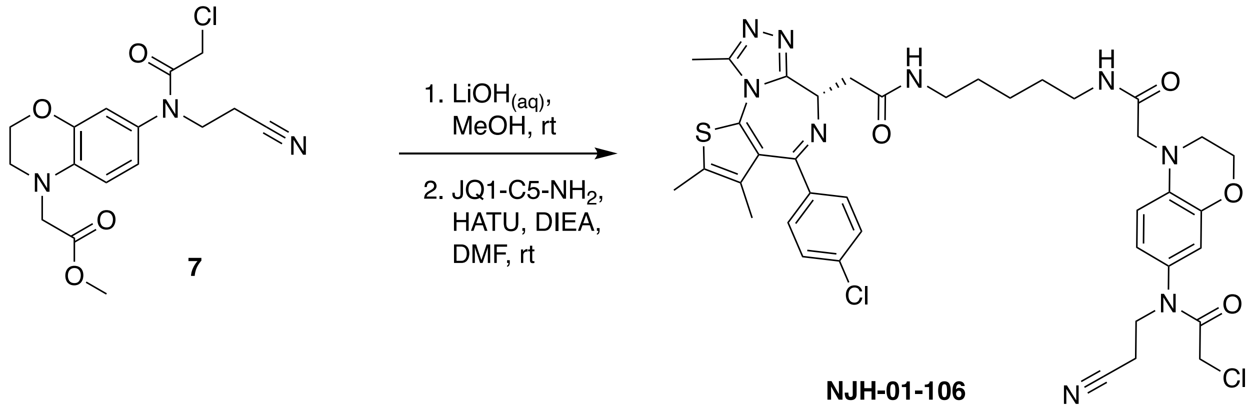


**NJH-01-106**

(S)-2-chloro-N-(4-(2-((5-(2-(4-(4-chlorophenyl)-2,3,9-trimethyl-6H-thieno[3,2-f][1,2,4]triazolo[4,3-a][1,4]diazepin-6-yl)acetamido)pentyl)amino)-2-oxoethyl)-3,4-dihydro-2H-benzo[b][1,4]oxazin-7-yl)-N-(2-cyanoethyl)acetamide (**NJH-01-106**): Chloroacetamide **6** (22 mg, 0.064 mmol) was dissolved in MeOH (600 mL) and treated with aqueous LiOH (400 mL, 0.5 M, 0.2 mmol) at rt for 1h. The mixture diluted with DCM (5 mL) and acidified with aqueous HCl (400 mL, 1 M) and the mixture extracted with DCM. Combined organic extracts were dried over $Na_2SO_4$, concentrated, and the crude residue dissolved in DMF (0.5 mL). DIEA (55 mL, 0.32 mmol) was added followed by **7** (31mg, 0.064 mmol) then HATU (49 mg, 0.13 mmol). The resulting solution was stirred for 10 minutes before it was diluted with EtOAc (0.5 mL) and purified by silica gel chromatography (0-10% MeOH/DCM) followed by preparatory thin layer chromatography (7% MeOH/DCM) to obtain the title compound (26 mg, 0.032 mmol, 50%) as a white lyophilized solid. HRMS $[M+H]^+$ calc 804.2535, found 804.2669. 1H NMR (400 MHz, CDCl3) δ 7.44 – 7.31 (m, 4H), 7.04 – 6.96 (m, 1H), 6.74 (dd, J = 8.5, 2.5 Hz, 2H), 6.68 (d, J = 2.5 Hz, 1H), 6.59 (d, J = 8.5 Hz, 1H), 4.62 (dd, J = 8.8, 5.3 Hz, 1H), 4.30 (t, J = 4.5 Hz, 2H), 3.89 (d, J = 5.5 Hz, 6H), 3.55 (dd, J = 14.4, 8.8 Hz, 1H), 3.46 (dq, J = 7.7, 3.8, 3.2 Hz, 1H), 3.43 – 3.32 (m, 1H), 3.32 – 3.24 (m, 1H), 3.23 – 3.13 (m, 2H), 2.69 (dd, J = 6.9, 2.9 Hz, 2H), 2.67 (s, 3H), 2.41 (d, J = 0.9 Hz, 3H), 1.70 – 1.65 (m, 3H), 1.55 – 1.41 (m, 6H), 1.35 (h, J = 6.0, 5.5 Hz, 2H). 13C NMR (151 MHz, CDCl3) δ 170.49, 169.20, 167.15, 164.08, 155.65, 149.90, 145.04, 136.92, 136.55, 135.77, 132.05, 131.07, 130.96, 130.94, 130.48, 129.85, 128.78, 121.05, 117.68, 115.75, 113.37, 64.79, 56.08, 55.58, 54.53, 48.69, 46.11, 41.91, 39.32, 39.15, 39.04, 28.70, 23.78, 18.65, 17.30, 16.28, 14.38, 13.11, 11.81.

NMR Spectra
NJH-2-030 1H and 13C NMR:

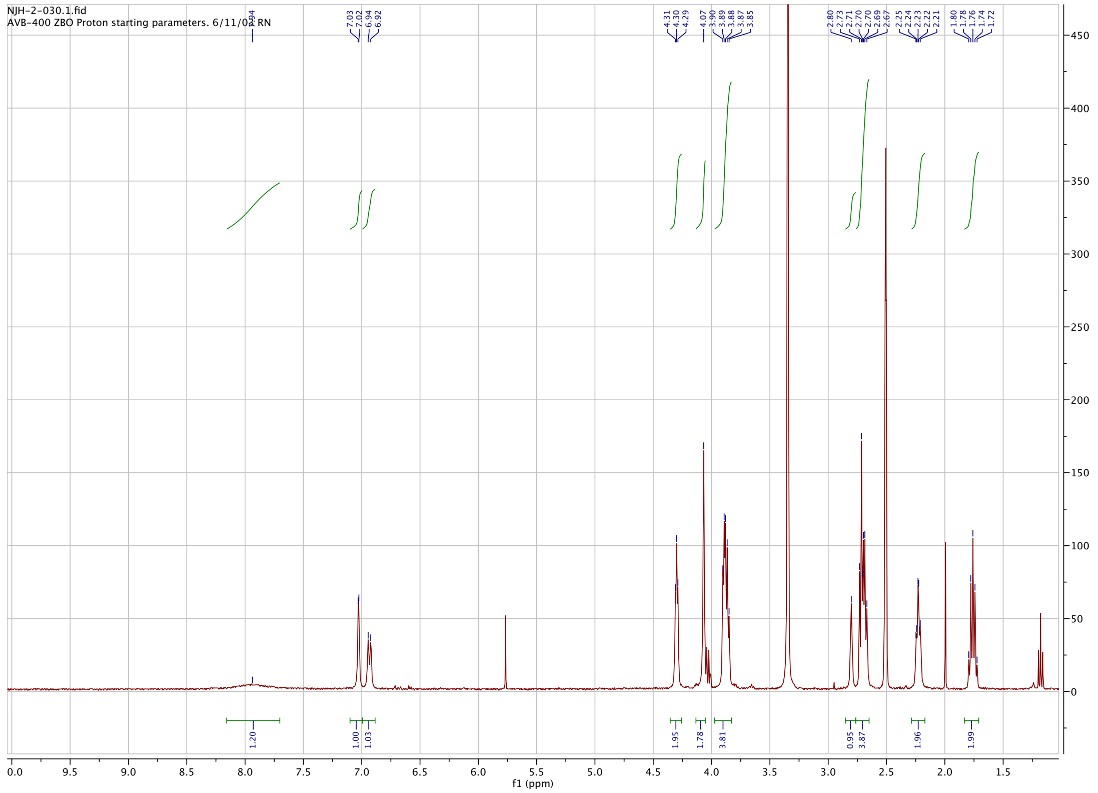


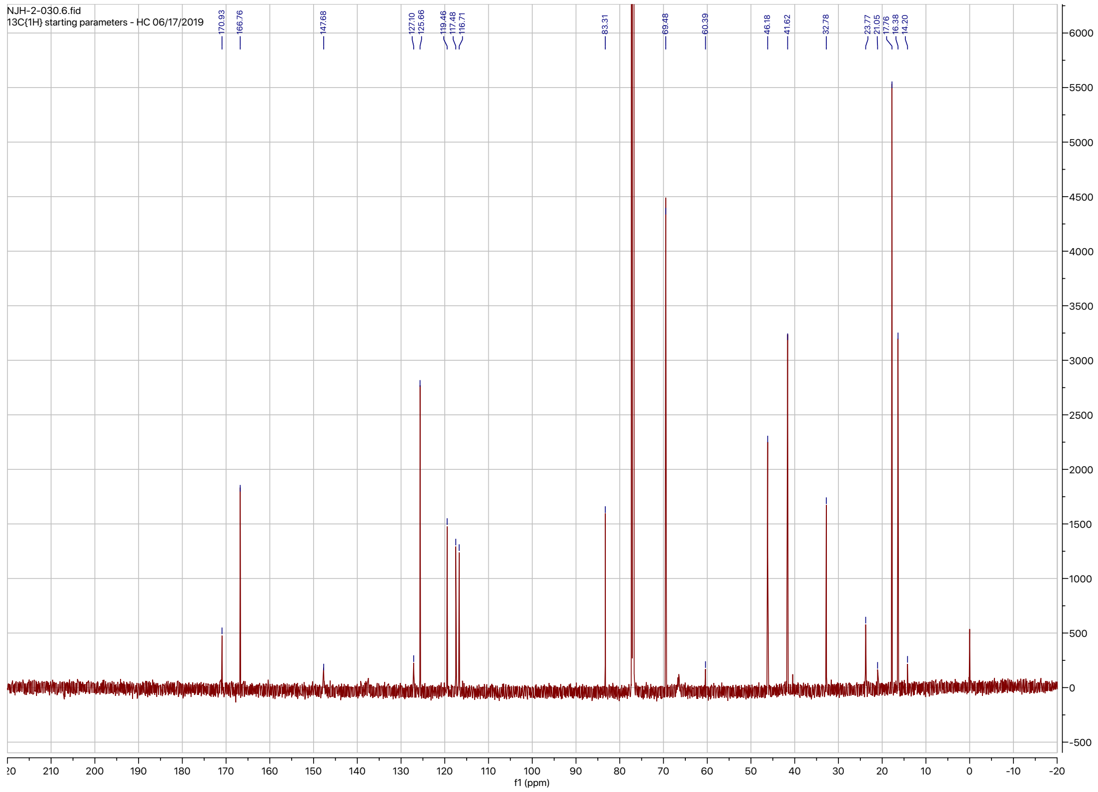

### NJH-01-106 1H and 13C NMR:

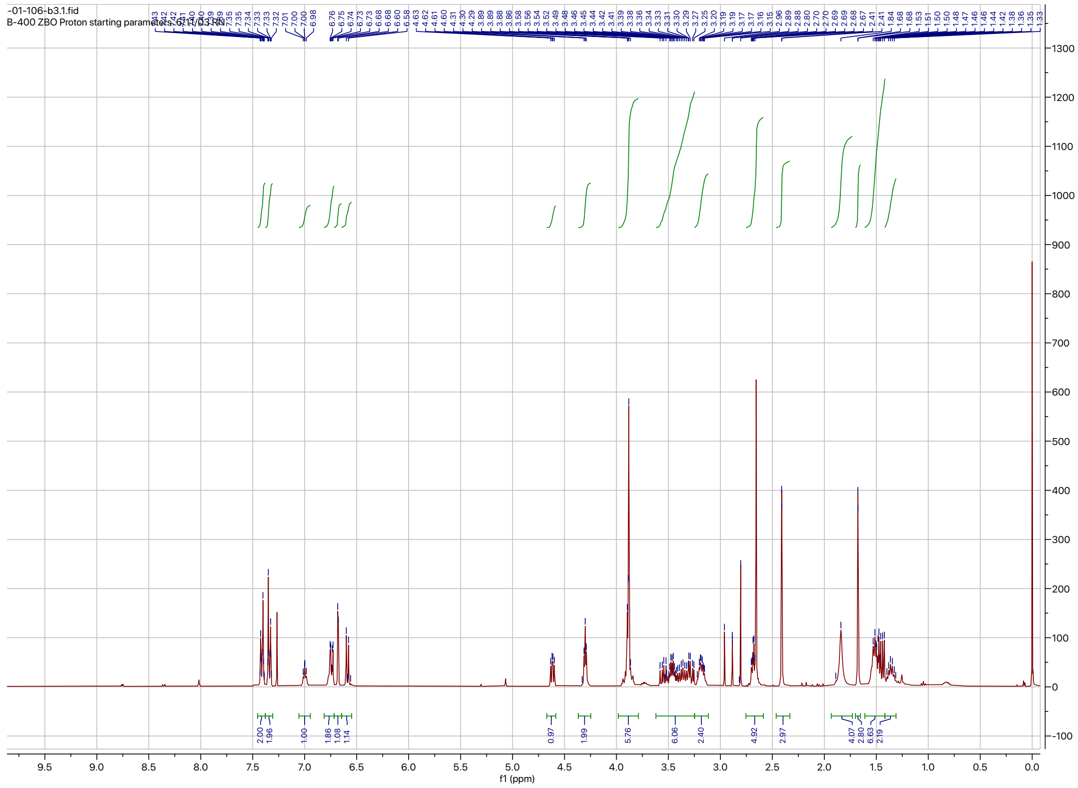

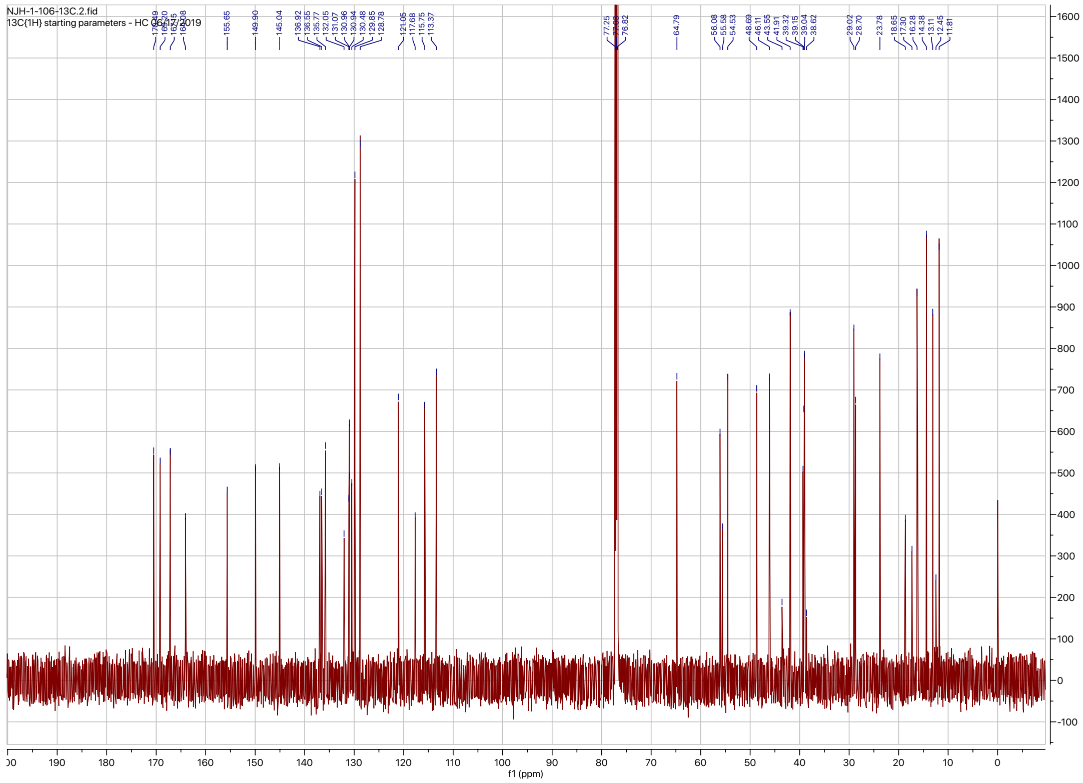

**Supporting Table Legends**

**Table S1. Cysteine-reactive covalent ligand screening against FEM1B. Tab 1** shows the compound name, Enamine catalog number, molecular weight, and structures of the compounds screened. **Tab 2** shows data from screening a cysteine-reactive covalent ligand library in a fluorescence polarization assay with TAMRA-conjugated FNIP1$^{562\text{-}591}$ degron with recombinant MBP-tagged FEM1B. FEM1B was pre-incubated with DMSO vehicle or covalent ligand (50 μM) for 30 min prior to addition of the TAMRA-conjugated degron.

**Table S2. Proteomic profiling of NJH-01-106 treatment in HEK293T cells.** HEK293T cells were treated with DMSO vehicle or NJH-01-106 (1 μM) for 12 h. Protein level changes in cell lysate were quantitatively assessed by TMT-based proteomic profiling. The first tab shows all identified proteins across three biologically independent replicates/group. The second tab shows quantified proteins with at least 2 unique peptides.

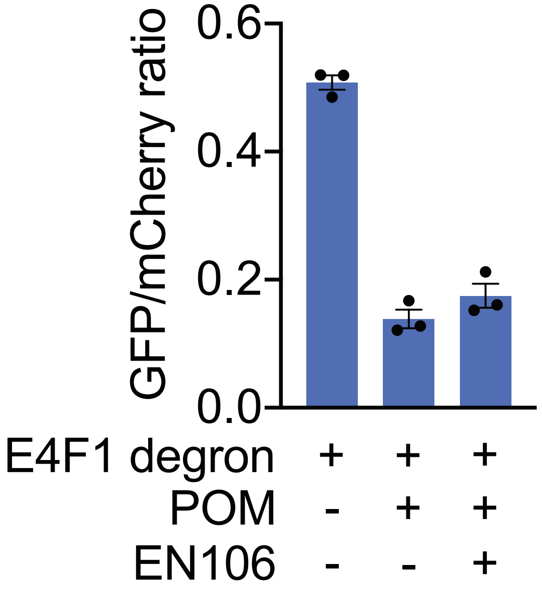


**Figure S1.** Flow cytometry analysis of E4F1-GFP degron levels compared to mCherry levels with DMSO vehicle, pomalidomide (10μM, 4h), or pomalidomide (10μM, 4h) and EN106 (10μM, 12h) treatment in HEK293T cells.
